# Common Variant Associations with Fragile X Syndrome

**DOI:** 10.1101/115998

**Authors:** James J Crowley, Jin Szatkiewicz, Anna K Kähler, Paola Giusti-Rodriguez, NaEshia Ancalade, Jessica K Booker, Jennifer L Carr, Greg E Crawford, Molly Losh, Craig A Stockmeier, Annette K Taylor, Joseph Piven, Patrick F Sullivan

## Abstract

Fragile X syndrome is a common cause of intellectual disability. It is usually caused by a de novo mutation which often occur on multiple haplotypes and should not be detectible using genome-wide association (GWA). We conducted GWA 89 male FXS cases and 266 male controls, and detected multiple genome-wide significant signals near FMR1 (odds ratio=8.10, P=2.5×10^−10^). These findings withstood robust attempts at falsification. Fine-mapping did not serve to narrow the interval (minimum P=1.13×l0^−14^), and functional genomic integration (including 5C data we generated for this region) did not provide a mechanistic hypothesis. Controls carrying a risk haplotype had significantly longer and more variable FMR1 CGG repeats than controls with the protective haplotype (P=4.75×10^−5^) which may predispose toward increases in CGG number to the pre-mutation range over many generations. This is a salutary reminder of the complexity of even “simple” monogenetic disorders.

## Introduction

Fragile X syndrome (FXS) (1) is a common cause of intellectual disability (0.25−1/1,000 male births) (2, 3). It is characterized by intellectual disability, autistic behavior, hyperactivity, anxiety, and pleomorphic physical abnormalities (e.g., tall stature, macroorchidism). (4) FXS is caused by CGG expansion in the 5’ UTR of the chromosome X gene *FMR1* in most cases. (5-7) Full FXS mutations are characterized by expansion of the *FMR1* 5’ UTR CGG repeat to ≥200 copies with pre-mutations in the 55-200 copy range. (8)

*FMR1* 5’ UTR CGG expansions generally arise as *de novo* mutations when mutable pre-mutations expand to full mutations during oogenesis. Although the probability of *de novo* mutations can be influenced by local DNA features, detection of *de novo* events using linkage disequilibrium would be unexpected for high-penetrance single gene disorders. (9, 10) This implies that genome-wide association (GWA) of FXS cases versus controls should not detect the *FRM1* region as a susceptibility locus for FXS. As part of a study of FXS and autism, we conducted a case-control GWAS for FXS.

## Materials and Methods

### Subjects

Males with a genetically-confirmed diagnosis of FXS were recruited from volunteer registries (URLs). All available medical records were reviewed, and any features suggestive of a complex or atypical presentation led to exclusion. Controls were males from the Genes and Blood Clotting Study (GABC) in dbGaP (URLs, accession phs000304.vl.pl). GABC participants were male university students who volunteered for a study of the genetics of hemostasis and who had no acute or chronic illnesses. Additional male comparison subjects were from the Swedish Schizophrenia Study (N=3,525, 46.4% cases). (11) As there is no evidence for association with schizophrenia in the *FMR1* region, (11, 12) cases and controls were combined. We also used male HapMap3 founders from northwestern Europe and Tuscany (CEU and TSI, N=101). (13) All procedures were approved by Institutional Review Boards, and written informed consent was obtained from the parents/legal guardians of FXS cases and from control subjects.

### Genetic assays

Table 1 summarizes the samples and assays used in this study. *<u>FMR1</u>*: the number of CGG repeats in the 5*’*UTR of *FMR1* was determined in 89 FXS cases with a validated diagnostic assay (Kimball Genetics, Denver, CO). (14-16) To understand the internal structure of *FMR1* CGG repeats and to place these on common haplotypes, we used AmplideX *FMR1* PCR kits (Asuragen, Inc; Austin, TX; catalog #49402) to quantify *FMR1* 5’UTR repeat sizes and to count AGG interruptions. <u>GWAS:</u> FXS cases and GABC controls were genotyped with lllumina HumanOmnil-Quad arrays, and genotypes were called using predefined clusters using GenomeStudio. Quality control was performed using PLINK. (17) SNPs were excluded for missingness > 0.03, minor allele frequency (MAF) < 0.01, deviation from Hardy-Weinberg expectations in controls (*P****<*** 1×10^-6^), SNP missingness differences between cases and controls (*P* < 0.05), or if a SNP probe did not map uniquely to the human genome. Subjects were excluded for missingness > 0.05, excessive autosomal homozygosity or heterozygosity, or relatedness (π ̂> 0.2 based on LD pruned autosomal SNPs). One FXS case was genotyped in duplicate with 0.99998 concordance, and a CEPH sample previously assayed with the same array had 0.99981 concordance. <u>TaqMan:</u> rs2197706 and rs5905149 genotypes were verified with TaqMan Assays (catalog #4351379 and #4351379, Applied Biosystems, Carlsbad, CA). A SNP from Gerhardt et al. (18) (rs45631657) was genotyped with a custom TaqMan assay. <u>Sequenom:</u> we designed two massARRAY iPLEX (San Diego, CA) genotyping panels for common variant fine-mapping. SNPs were selected from GWAS results, haplotype analyses, and common variation databases, and then pruned using TAGGER. (19) All assays used genomic DNA isolated from peripheral blood. The genome reference was GRCh37/UCSC hgl9.

**Table 1.** Summary of samples and genotyping.

| Purpose | Genotyping | FXS Cases | Controls | Control source |
| --- | --- | --- | --- | --- |
| Establish FXS status | *FMR1* 5'UTR CGG repeat (CLIA assay) | 89 | 0 | N/A |
|  | *FMR1* CGG repeat analysis | 82 † | 109 | Sweden |
| GWAS | Illumina HumanOmni1-Quad | 89 | 266 | GABC |
| Allele frequency comparisons | Illumina Human1M & Affymetrix 6.0 | – | 101 | HapMap3 |
|  | Illumina OmniExpress | – | 3,525 | Sweden |
| Verify key genotypes | TaqMan rs2197706 | 82 † | 0 | N/A |
|  | TaqMan rs5905149 | 82 † | 94 | Sweden |
| Common variant fine-mapping | Sequenom (31 SNPs) | 97 | 467 | Sweden |
|  | TaqMan rs45631657 | 103 | 467 | Sweden |

† insufficient DNA for 7 FXS cases.

### Statistical analysis

Case-control comparisons were performed using PUNK (17) using logistic regression under an additive model with three ancestry principal components as covariates.

## Results

We conducted GWA analyses for 750K SNPs in 89 male FXS cases and 266 male controls **(*Table S1*).** All FXS cases had full *FMR1* mutations (>200 for the 5’UTR CGG repeat), and this was verified with a second assay (82 FXS cases with sufficient DNA). We assessed ancestry using principal components analysis (20) on LD-pruned autosomal SNPs **(*Figure S1*).** All controls and 90% of cases were of predominant European ancestry (we retained nine cases of mixed ancestry given the small number of cases). Logistic regression analyses identified five SNPs that met genome-wide significance with odds ratios > 5 **(*Table2, Figures 1a-b*).** These SNPs were in a 66 kb interval from chrX:146.85-146.92 Mb located 75 kb 5’ of the nearest gene, *FMR1*. Repeating the logistic regression conditioning on the most strongly associated SNP (rs2197706) markedly attenuated significance in the *FMR1* region suggesting the presence of a single association signal.

Given that FXS usually results from *de novo* mutations, strong associations with common variation are unexpected. Indeed, the strongest association (rs2197706, odds ratio=8.10, *P*=2.5×l0^−10^) is among the top dichotomous trait associations in the NHGRI/EBI GWAS catalog (21) (URLs). We therefore evaluated alternative explanations for these findings. First, given the marked allele frequency differences in cases and controls, re-genotyping rs5905149 and rs2197706 with TaqMan assays showed perfect agreement with Illumina array genotypes, and served to exclude allele assignment errors. Second, the allele frequencies in cases and controls were similar genome-wide except for SNPs 5’ of *FMR1* (***Figure S2*).** Given our use of controls genotyped independently from cases, it is important to note that the control allele frequencies for the significant SNPs in the *FMR1* region were similar to those from two external samples **(*Table 2*).** Third, genome-wide *P*-values conformed closely to the null expectation (mean *P*-value=0.500 over 750K SNPs, Figure 1a), findings inconsistent with uncontrolled bias. Fourth, exclusion of nine cases with mixed ancestry had little impact on the results **(*Table 2*)**. Fifth, a trivial explanation for these findings is if cases were cryptically related via a recent shared ancestor; however, case-case pairs were slightly <u>less</u> related on average than control-control pairs (***Figure S3***). Cases and controls had similar proportions of autosomal homozygous SNPs as well as the number and size of autosomal runs of homozygosity (no comparisons were significantly different and cases had lower means in each instance). Sixth, asymptotic *P*-values can be inaccurate in small samples, but Fisher’s exact test and permutation procedures yielded similar significance levels. Thus, we could identify no plausible alternative explanation for our findings.

**Figure 1.**
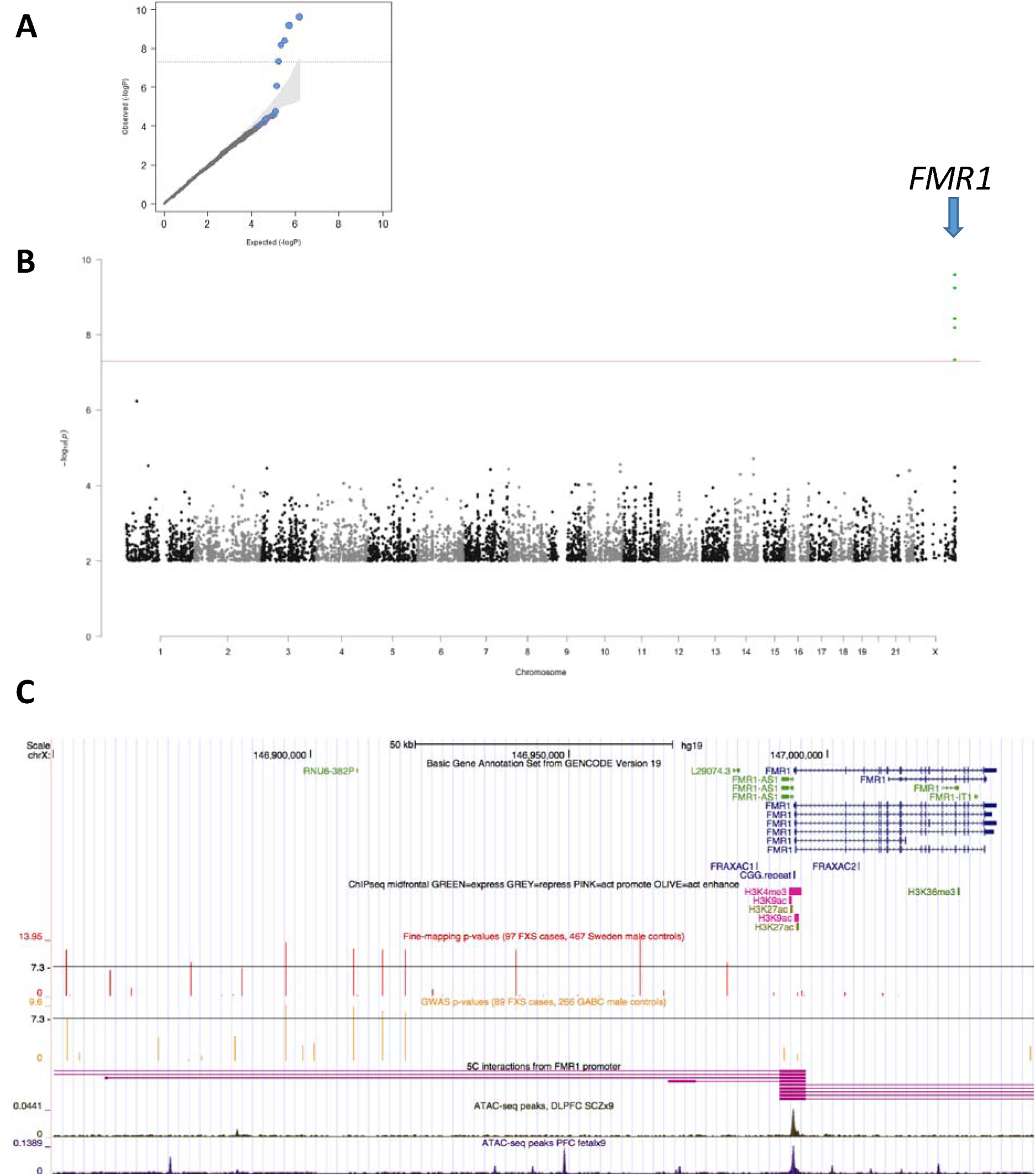
FXS case-control GWAS. (A) Quantile-quantile plot for logistic regression of male FXS cases and GABC controls (including ancestry principle components). The observed P-values conform closely to the null except for five SNPs in the FMR1 region. The shaded region indicates the expected 95% probability interval for ordered P-values. (B) Manhattan plot for the GWAS of male FXS cases and GABC controls (logistic regression including ancestry principle components). The X-axis is chromosomal position from 1ptel to Xqtel. The Y-axis is −log_10_(P). Genome-wide significant SNPs near FMR1 are indicated. (C) Detailed of FMR1 region (hg19, chrX:146850000-147040000). Tracks are: GENCODE gene annotations; positions of FRAXAC1, FRAXAC2, and promoter CGG repeat; selected ChIP-seq marks; SNP positions and −log_10_(P) for SNPs in the fine-mapping study and in the GWAS; DNA-DNA chromosomal looping from 5C based on the FMR1 promoter; and open chromatin in pre-frontal cortex of 9 adult SCZ cases and 9 fetal samples.

**Table 2.** Genome-wide significant results of GWAS of male FXS cases and controls.

| SNP | chrX (hg19) | Alleles | OR (95% CI) | *P* | $F_{\text{case}}$ | $F_{\text{control}}$ | $F_{\text{Sweden}}$ | $F_{\text{CEU}}$ | $F_{\text{TSI}}$ |
| --- | --- | --- | --- | --- | --- | --- | --- | --- | --- |
| rs5952060 | 146852679 | C/T | 5.33 (2.93-9.71) | $4.57 \times 10^{-8}$ | 0.764 | 0.365 | 0.357 | 0.386 | 0.409 |
| rs2197706 | 146895120 | A/C | 8.10 (4.24-15.49) | $2.53 \times 10^{-10}$ | 0.807 | 0.351 | – | 0.357 | 0.386 |
| rs5905149 | 146908213 | A/C | 5.99 (3.40-10.56) | $5.76 \times 10^{-10}$ | 0.584 | 0.184 | 0.183 | – | – |
| rs7876251 | 146913828 | G/A | 5.64 (3.17-10.01) | $3.68 \times 10^{-9}$ | 0.693 | 0.286 | 0.279 | 0.263 | 0.279 |
| rs4824253 | 146918268 | G/A | 5.35 (3.04-9.43) | $6.41 \times 10^{-9}$ | 0.685 | 0.286 | 0.278 | 0.263 | 0.296 |

The first allele given is the least common in this sample and is the reference for the odds ratio (OR) and frequencies. CI is confidence interval. *P* is from the logistic regression including ancestry covariates. Logistic regression *P*-values after removing nine cases with divergent ancestry dropping were $5.86 \times 10^{-8}$ , $3.36 \times 10^{-10}$ , $8.52 \times 10^{-10}$ , $4.79 \times 10^{-9}$ , and $8.38 \times 10^{-9}$ (respectively). Shown are allele frequencies for male FXS cases ( $F_{\text{case}}$ ), GABC controls ( $F_{\text{control}}$ ), subjects from Sweden ( $F_{\text{Sweden}}$ ), and HapMap3 northwestern European ( $F_{\text{CEU}}$ ) and Tuscan control samples ( $F_{\text{TSI}}$ ).

In a fine-mapping experiment, we genotyped 32 SNPs (chrX: 146844358-147013704, the association region extending into *FMR1*) in an expanded set of 97 FXS cases and 467 male controls (from a different study than for the initial GWAS). We included rs45631657which was reported to inactivate an important replication origin. (18) Variable numbers of SNPs overlapped with independent genotypes on the same subjects, and we observed 100% agreement (data not shown). Nine SNPs exceeded genome-wide significance **(*Table S2*).** All five SNPs in Table 2 replicated with consistent odd ratios and greater significance (*P*-values ranging from 4×10^−12^ to 7×10^−14^), and four other SNPs reached genome-wide significance (rs4824231, *P*=1.13×10^−14^; rs25705, P=5.74×10^−9^; rs45631657, *P*=5.20×10^−12^; and rs112146098, *P*=6.60×10^−9^). Repeating the logistic regression conditioning on rs2197706 or rs4824231 markedly attenuated significance in the *FMR1* region suggesting the presence of a single association signal. Thus, we continued to observe a broad region of significance.

Figure 1c depicts the 128 kb association region, from 141 kb to 13 kb 5’ of *FMR1*. The association region includes the *FMR1* promoter CGG repeat. Table 3 shows haplotypes from the genome-wide significant SNPs. The most common haplotype was strongly protective, and there were two risk haplotypes. *FRM1* CGG analysis was available on 81 controls. Controls with the risk haplotype had significantly longer CGG repeats than controls with the protective haplotype (median 15 and interquartile range 13-22 versus median 10 and interquartile range 10-10, *F_1,79_=* 18.3, *P*=4.75 × 10^−5^). There was greater variability in CGG number in controls with the risk that the protective haplotype (standard deviation 2.4 vs 6.8). Around 40% of cases had additional phenotype measures (e.g., Vineland Adaptive Behavior Scale and the Social Responsiveness Scale) and there were no significant differences between cases with the risk or protective haplotypes (data not shown).

**Table 3.** Haplotype analyses of FMR1 region.

| Haplotype | Subjects | Controls | FXS cases | Freq control | Freq case |
| --- | --- | --- | --- | --- | --- |
| TGACGGTCC | 17 | 14 | 3 | 0.030 | 0.031 |
| CGACGGTCC | 20 | 17 | 3 | 0.036 | 0.031 |
| CGAAGGTTT | 21 | 20 | 1 | 0.043 | 0.010 |
| CGACAATTC | 26 | 17 | 9 | 0.036 | 0.093 |
| CGCCAATCC | 31 | 29 | 2 | 0.062 | 0.021 |
| CGAAGGTTC | 65 | 42 | 23 | 0.090 | 0.237 |
| CCAAGGCTT | 77 | 44 | 33 | 0.094 | 0.340 |
| TGCCAATCC | 261 | 250 | 11 | 0.535 | 0.113 |

Observed haplotypes from the fine-mapping data (32 SNPs in 97 FXS cases and 467 male controls). Haplotypes were created using nine genome-wide significant SNPs (*rs5952060-rs112146098-rs2197706-rs5905149-rs7876251-rs4824253-rs45631657-rs4824231-rs25705*). Haplotypes counts <10 (N=46) were removed. Logistic regression highlighted a strongly protective haplotypes (green) and two risk haplotypes (red).

We next evaluated possible functions of the association region using functional genomic data **(*Figure 1c*)**. Using RNA-seq data from human dorsolateral prefrontal cortex (DLPFC) in nine SCZ cases and nine controls along with prefrontal cortex (PFC) from nine fetuses and three neural progenitor cell lines, we saw that *FMR1* (but not the antisense transcript, *FMR1-AS1)* was robustly expressed. There was no evidence of substantial gene expression or an unannotated feature in the association region 5’ to *FMR1*. The expression of *FMR1* in DLPFC is associated with a common genetic variant but the associated SNP is far outside the region. (22) We evaluated DNA-DNA interactions using 5C (chromosome conformation capture-carbon copy) from five human fetal prefrontal cortex samples for a 1.26 Mb region overlapping *FMR1* and the association region ***(Figure S5)*.** *FMR1* promoters had evidence of DNA-DNA interactions with the association region and with other parts of *FMR1*. A microRNA gene cluster around 700 kb 5’ to *FMR1* had more substantial DNA-DNA interactions ***(Figure S5)*** but had no genetic association signal. We identified open chromatin using ATAC-seq (human DLPFC in nine SCZ cases and nine controls along with PFC from nine fetuses). The SNPs associated with FXS were not notable for open chromatin or key ChIP-seq marks.

## Discussion

GWA of FXS cases versus controls identified an unusually strong association with the *FMR1* region. The largest association (odds ratio=8.10, *P*=2.5 ×10^−10^) is among the top dichotomous trait associations in the NHGRI/EBI GWAS catalog (21) (URLs), and generally exceeded only by rare adverse drug reactions. Given the small sample size (89 FXS cases and 266 male controls), it is notable that the association survived robust attempts at falsification.

In some respects, our identification of the causal locus for FXS - a rare, single-gene disorder - in an outbred population using a linkage disequilibrium-based approach like GWA is unexpected. GWA in case-control samples can detect rare causal genes in special circumstances that do not apply here (e.g., when cases inherit a causal mutation from a relatively recent common ancestor (23, 24) or if multiple rare mutations yield an aggregate signal detectible by GWA (25)). *De novo* mutations in particular may be invisible to GWA: although *de novo* mutational processes can be influenced by local genomic context, replication timing, and genotypes at other loci, (9,10, 26-28) these effects are generally not deterministic, and most *de novo* mutations occur on different haplotypes.

With the exception of unusual exonic mutations, FXS is caused by a *de novo* mutational event in the expansion of a pre-mutation to a full mutation during oogenesis. (5-7) However, the local genomic context of *FMR1 de novo* promoter mutations is influential. (29-31) There is substantial evidence that this region is detectible via linkage disequilibrium in case-control studies using a few microsatellite markers. (32) Indeed, a 1992 paper (33) reported a FXS case versus control haplotype difference as "P<0.001" but the P-value actually reached genome-wide significance (P ~9x10^−9^). The association of common variation upstream of *FMR1* with FXS has strong replication evidence in the literature: this is unquestionably a true association.

Fine-mapping of the interval and integration with a number of types of functional genomic data did not narrow the region or yield a mechanistic hypothesis. A lack of early fetal data limits this conclusion. It is possible that a population genetic mechanism is at work: the risk haplotype is present in ~18% of European-ancestry controls, and tends to carry a greater and more variable number of CGG repeats which may predispose toward increases in CGG number to the pre-mutation range over many generations. A similar mechanism has been reported for Huntington’s disease. (34) This is a salutary reminder of the complexity of even “simple” monogenetic disorders.

## Conflicts of Interest

PFS is a scientific advisor for Pfizer, Inc. and received an honorarium from F. Hoffmann-La Roche AG.

## Acknowledgements

This project was funded by an Autism Speaks (URLs) award to PFS. PFS gratefully acknowledges support from the Swedish Research Council (Vetenskapsrådet, award D0886501). We are indebted to Dr Mark Daly for discussions regarding the results, and to Dr Job Dekker and his group for assistance with the 5C work. For the human postmortem samples, the authors acknowledge the Cuyahoga County Medical Examiner’s Office and the families of the deceased. They also note contributions of Drs. James Overholser and George Jurjus and of Lesa Dieter in the retrospective psychiatric assessments, and Dr. Grazyna Rajkowska and Gouri Mahajan in sample preparation - this work was supported by NIH/NIGMS COBRE Center for Psychiatric Neuroscience (P30 GM103328).

## URLs

Fragile X Research Registry, https://www.fragilexregistrv.org. dbGaP,http://www.ncbi.nlm.nih.gov/gap. NHGRI GWAS catalog, https://www.ebi.ac.uk/gwas. Autism Speaks, http://www.autismspeaks.org

